# Structural connectome architecture shapes the maturation of cortical morphology from childhood to adolescence

**DOI:** 10.1101/2022.12.15.520527

**Authors:** Xinyuan Liang, Lianglong Sun, Xuhong Liao, Tianyuan Lei, Mingrui Xia, Dingna Duan, Zilong Zeng, Qiongling Li, Zhilei Xu, Weiwei Men, Yanpei Wang, Shuping Tan, Jia-Hong Gao, Shaozheng Qin, Sha Tao, Qi Dong, Tengda Zhao, Yong He

## Abstract

Cortical thinning is an important hallmark of the maturation of brain morphology during childhood and adolescence. However, the connectome-based wiring mechanism that underlies cortical maturation remains unclear. Using neuroimaging, connectome, transcriptome, and computational modeling, we mapped cortical thinning patterns primarily located in lateral frontal and parietal heteromodal nodes during childhood and adolescence, which is structurally constrained by white matter network architecture and is particularly represented using a network-based diffusion model. Furthermore, connectome-based constraints are regionally heterogeneous, with the largest constraints residing in frontoparietal nodes, and are associated with gene expression signatures of microstructural neurodevelopmental events. These results are highly reproducible while using another independent dataset. Our findings advance our understanding of network-level mechanisms and the associated genetic basis that underlies the maturational process of cortical morphology during childhood and adolescence.

## MAIN TEXT

### Introduction

The transition period from childhood to adolescence is characterized by prominent reorganization in cortical morphology (*1, 2*), which provides critical support for cognitive growth (*3, 4*). With progress in modern *in vivo* structural brain imaging, researchers have documented widespread spatial refinements of cortical morphology throughout childhood to adolescence (*5, 6*). A typical cortical maturation sequence is marked by hierarchical cortical thinning from the primary to association cortex (*1, 7, 8*) and is thought to be mediated by cellular mechanisms, genetic regulation, and biomechanical factors (*9, 10*). In the present study, we present a mechanistic approach to model how the maturational pattern of cortical morphology from childhood to adolescence is shaped by white matter (WM) connectome architecture.

At the microscale level, a large quantity of histological studies has suggested that the brain’s WM pathways are involved in the developmental process of cortical gray matter. During neural circuit formation, axons express guidance receptors to integrate attractive and repulsive environmental information for navigation to their target neurons (*11, 12*). After axons arrive, synaptic maintenance and plasticity rely on active axonal transport through axonal cytoskeletons, which offers essential delivery of neurotrophic factors, energy requirements, and synthesized or degraded proteins for long distanced cortical neurons (*13–15*). Such early-established neuronal pathways could lead to preferences in attracting or removing new links during the formation of cortical hubs (*16*). Physical simulation studies suggest that there may be a tension-induced relationship between fiber growth and cortical fold morphology (*17, 18*). At a macroscale level, several prior studies using structural and diffusion MRI have also shown that focal cortical thickness (CT) decreases are associated with increased microstructural anisotropy and decreased mean diffusivity in adjacent WM (*19–22*) and that homologous cortical regions, which are rich in WM fibers, exhibit higher maturational couplings of CT than nonhomologous regions (*21, 23*).

Notably, all these previous studies are limited to local cortical regions or certain fiber tracts. The human brain is a highly interacting network in nature in which connections promote interregional communications, raising the possibility that the maturation of focal cortical morphology is shaped by the overall architecture of the WM connectome. However, whether and how the maturation pattern of cortical morphology from childhood to adolescence is constrained by physical network structures, and specifically, whether this constraint works following a network-based diffusion model, remains largely unknown. We anticipate that models of regional cortical maturation would yield mechanistic insights into network structure that govern the coordinated development of cortical morphology among regions.

If the connectome structure shapes regional cortical maturation, it is necessary to further clarify whether this constraint is regulated by genetic factors. Converging evidence indicates that genetic modulations may exist on the potential constraint of WM maturation on cortical morphology. Studies on the rodent nervous system (*24, 25*) have shown that the wiring diagram of brains is tracked by genes that are involved in axon guidance and neuronal development processes. Such genetic cues are also related to molecules responsible for cytoskeletal rearrangements that induce cortical refinement processes, including synaptic pruning and neuron cell death (*12*). In humans, recent emerging transcriptome imaging analyses pave a new way to link brain macroscale structural maturation to microscale biological processes by seeking linkage between MRI-based brain measurements and genetic samples of postmortem brains. Such studies have shown that cortical thinning during maturation is related to genes involved in the structure and function of synapses, dendrites, and myelin (*26, 27*). These precisely programmed microstructural alterations constitute major neurodevelopmental events that promote the establishment of more mature brain architecture and anatomical connectivity from childhood to adolescence (*28, 29*). Therefore, we further hypothesize that the constraint between the maturation of cortical morphology and WM network structure is associated with gene expression profiles that are involved in neurodevelopment.

To fill these gaps, in the present study, we integrated neuroimaging, connectome, and transcriptome analyses as well as computational modeling to investigate network-level mechanisms underlying regional changes of cortical morphology during childhood and adolescence and to further explore their potential genetic underpinnings. Specifically, we test three hypotheses: (*i*) that the maturation of CT of brain nodes is associated with that of structurally connected neighbors, (*ii*) that the network-level diffusion model, which represents the direct and high-order information exchange preferences among neighbors, captures the principle of connectome constraint on the maturation of CT, and (*iii*) that the connectome constraints on cortical maturation are linked with gene expression levels of neurodevelopment processes.

## Results

### Data Samples

To investigate the relationship between cortical morphology maturation and the WM connectome from childhood to adolescence, we leveraged structural and diffusion MRI data from a longitudinal MRI dataset (“Discovery dataset”) with 521 brain scans from 314 participants (aged 6-14 years) in the Children School Functions and Brain Development Project in China (Beijing Cohort) (Fig. 1A). To obtain typical sample representations of childhood and adolescence phases, we divided all participants into the child group (218 participants, 299 scans, 6.08-9.98 y) and the adolescent group (162 participants, 222 scans, 10.00-13.99 y) using age 10 years as a cutoff, according to the criteria from a previous public health investigation (*30*) and the World Health Organization (WHO) (*31*). Besides this group-based analysis, we also validated our results by either considering age as a continuous variable or performing individual-based analysis. To assess the reproducibility of our results, we included an independent dataset (“Replication dataset”) that contains cross-sectional structural and diffusion MRI data of 301 typically developing participants, which were divided into the child group (98 participants, 5.58-9.92 y) and in the adolescent group (203 participants, 10.00-14.00 y) from the Lifespan Human Connectome Project in Development (HCP-D) (*32*). Details of the demographic information for all participants, data acquisition, and data analysis are provided in the Supplementary Text, Section 1.1-1.3.

**Figure 1.**
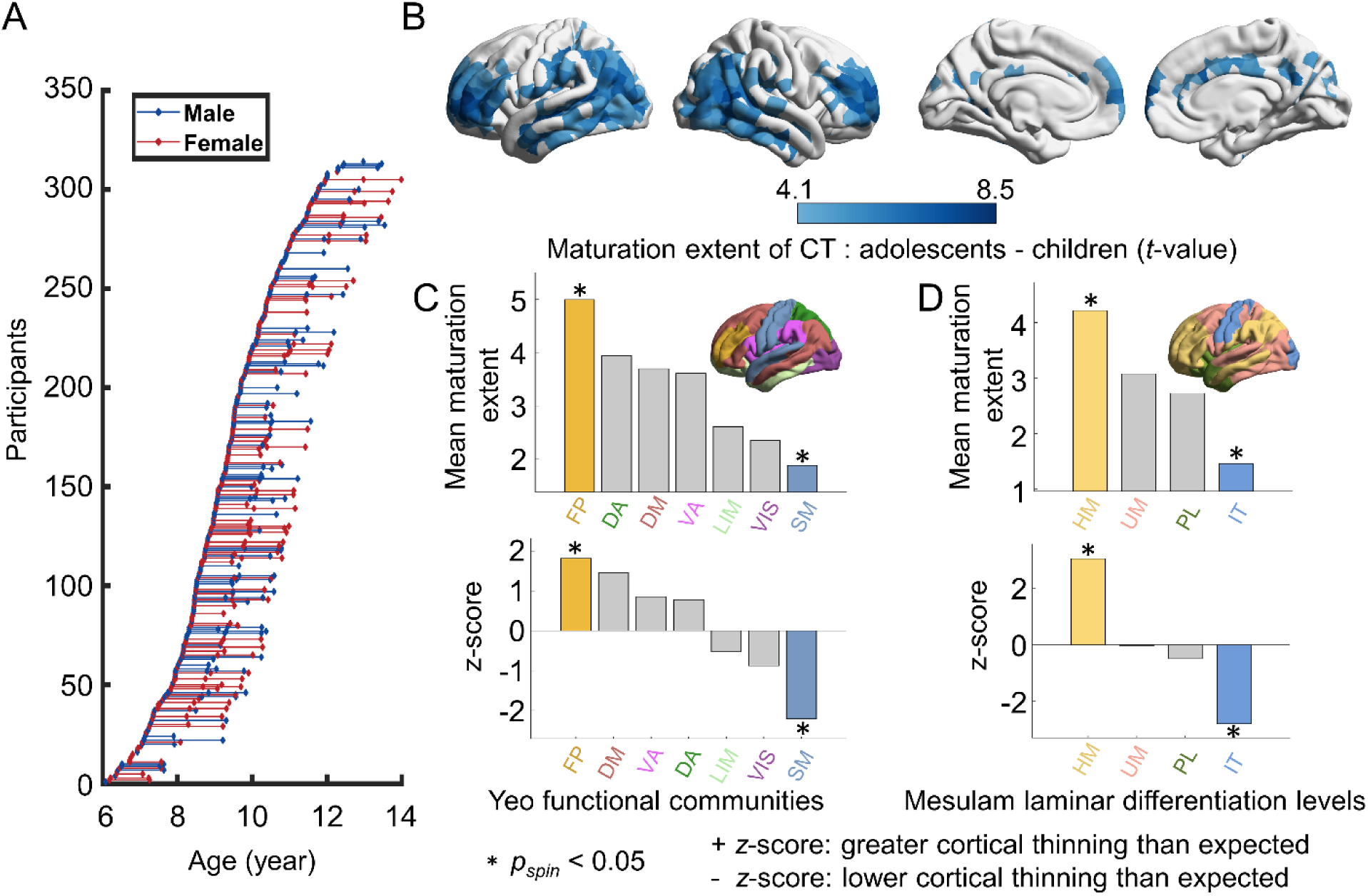
Scan age distribution of all participants and the typical CT maturation from childhood to adolescence. **(A)** Participants’ age at MRI scans. Each dot represents a single scan of a participant and the longitudinal scans of the same participant are connected by lines. (**B**) Statistical map of CT differences between child and adolescent groups. A greater positive *t*-value denotes more pronounced cortical thinning with development. The map was corrected using a Bonferroni correction method for multiple comparisons (*p_bonf_* < 0.05). (**C**) The mean CT maturation extent (estimated by *t*-value) within each brain community was defined by Yeo et al. (*37*), and the laminar differentiation level was defined by Mesulam et al. (*40*). (**D**) Spin tests (*38, 39*) were performed by spherical projection and rotation class positions 1000 times for correcting spatial autocorrelations, and the class-specific mean *t-*values were expressed as *z* scores relative to this null model. A positive *z* score indicated higher cortical thinning than expected by chance. Asterisks denote statistical significance (*p_spin_* < 0.05). VIS, visual; SM, somatomotor; LIM, limbic; DA, dorsal attention; VA, ventral attention; FP, frontoparietal; DM, default mode; IT, idiotypic; PL, paralimbic; UM, unimodal and HM, heteromodal. Values of a brain map were visualized using BrainNet Viewer (*112*).

### The Typical Spatial Refinement of Brain CT from Childhood to Adolescence

For each individual, we first parcellated the brain cortex into 1000 nodes of interest with approximately equal size (219 and 448 nodes parcellations as a validation (*33*) according to the modified Desikan-Kiliany atlas (*34, 35*). Then, we computed the average CT for each brain node based on structural MR images. To delineate the spatial maturation map of brain morphology, we estimated the statistical differences in regional CT between the child and adolescent groups to represent the CT maturation extent during development by a mixed linear analysis (*36*) with sex included as the covariate (Supplementary Text, Section 1.4). Brain nodes showed significant cortical thinning, mainly concentrated in dorsolateral prefrontal regions, lateral temporal and lateral parietal regions (Fig. 1B, *t*-values > 4.10, *P* < 0.05, Bonferroni corrected). To test whether this maturation pattern is anchored to specific brain systems, we classified all cortical nodes into seven well-validated brain communities (*37*) and performed a spherical projection null test (“spin test”) to correct for spatial autocorrelations by permuting seven communities positions 1000 times (*38, 39*). The class-specific mean *t*-values were expressed as *z* scores relative to this null model. We found that all brain systems showed decreased CT with development on average. The frontoparietal (FP) system and default mode (DM) system showed higher cortical thinning than expected by chance (FP: *p_spin_* = 0.029; DM: *p_spin_* = 0.068, Fig. 1C). The somatomotor (SM) system displayed lower cortical thinning than expected by chance (*p_spin_* = 0.004). We also repeated this analysis by classifying cortical nodes into four laminar differentiation levels (*40*). We found that heteromodal areas displayed cortical thinning (*p_spin_* < 0.001), while idiotypic areas showed lower cortical thinning than expected by chance (*p_spin_* = 0.001, Fig. 1D). Consistent results were found at the other two parcellation resolutions (Figs. S1-S2). These results are largely compatible with previous studies (*1, 8*), demonstrating that CT exhibits the most pronounced thinning in high-order association areas and is relatively preserved in primary areas from childhood to adolescence.

### Direct WM Connections Constrain the Spatial Maturation of CT

Next, we tested whether the regional maturation of CT was constrained by its direct WM connections. To this end, we first reconstructed individual structural connectomes with 1000 nodes (219 and 448 nodes as a validation) based on diffusion MR images of the child group by performing deterministic tractography between cortical regions (*41, 42*). We then generated a binary, group-level connectome by using a consensus approach that preserves the connection length distributions of individual networks (*43*) (Fig. 2A, Supplementary Text, Section 2.1).

**Figure 2.**
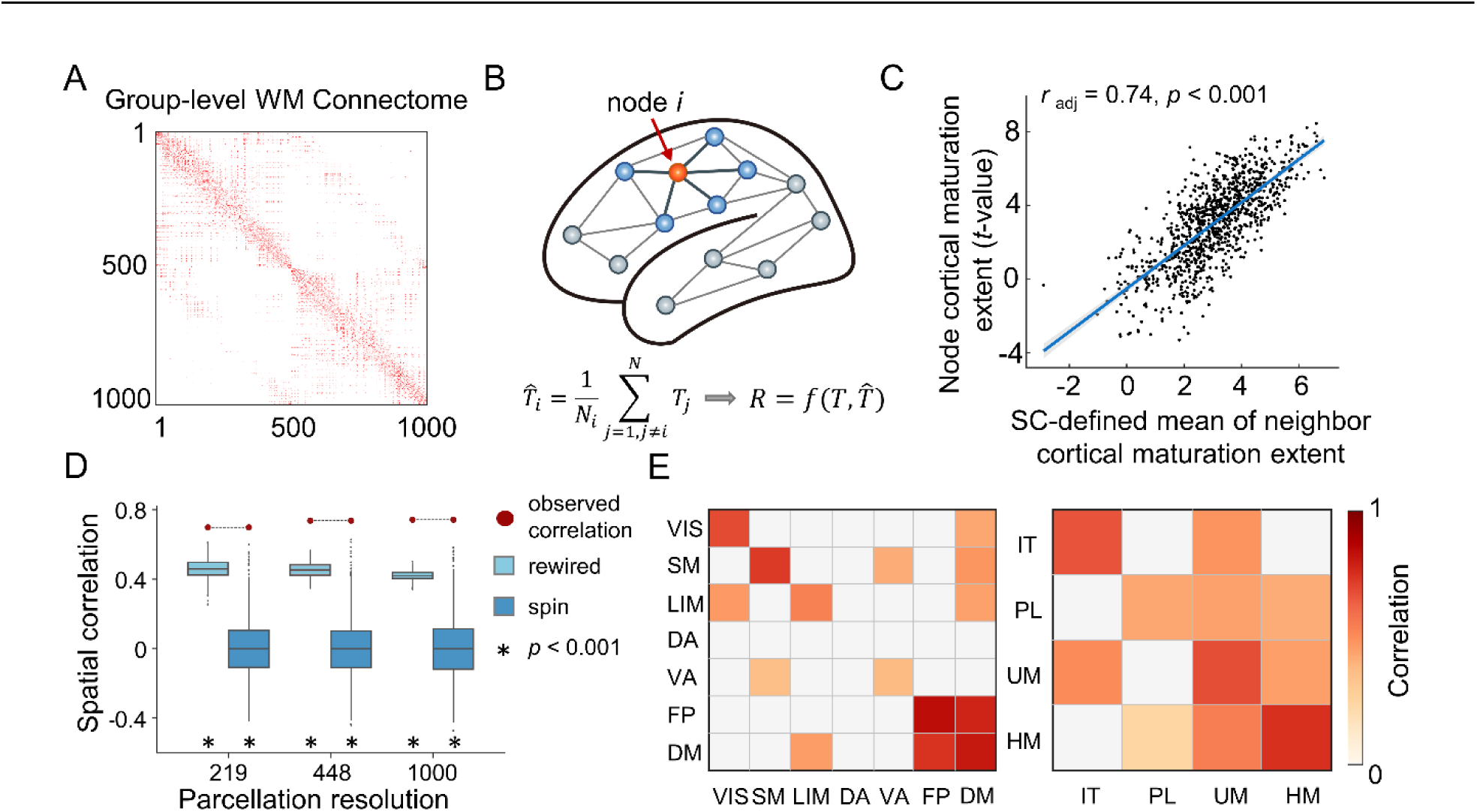
WM network-based CT maturation. (**A**) Group-level connectome backbone at 1000-node resolution. (**B**) Schematic diagram of WM network-constrained CT maturation. The CT maturation extent of a given node (orange) was correlated with the mean maturation extent of its directly connected neighbors (blue) to test whether the maturation of CT is shaped by the underlying WM network architecture. (**C**) A significant correlation was observed between the nodal CT maturation extent and the mean of its directly connected neighbors (*r_adj_* = 0.74, *P* = 5.56 × 10 ^-176^). The scatter plot shows the result at 1000-node resolution. See Fig. S3 for results at other resolutions. (**D**) The observed correlations across 3 resolutions (shown as red circles) were compared against two baseline null models. (1) To determine whether these correlations were driven by the basic spatial embedding of the WM network, we randomly rewired edges while preserving the nodal degree and edge length distribution of the empirical WM network (“rewired”, 1000 times, shown as light blue boxes). (2) To determine whether these correlations were driven by spatial autocorrelation, we generated 1000 surrogate maps by rotating region-level cortical *t* values (“spin test”, shown as deep blue boxes). Asterisks denote statistical significance (*p* < 0.001). (**E**) The spatial correlation at the system level. The whole-brain cortical nodes were classified into seven classic communities (*37*) (left) and four laminar differentiation levels (*40*) (right), and the statistically significant (*p_spin_* < 0.05, FDR corrected) correlations are shown in color. VIS, visual; SM, somatomotor; LIM, limbic; DA, dorsal attention; VA, ventral attention; FP, frontoparietal; DM, default mode; IT, idiotypic; PL, paralimbic; UM, unimodal and HM, heteromodal.

Next, we estimated the across-node relationship of the CT maturation extent (*t*-value between child and adolescent groups) between a node and its directly connected neighbor nodes in the backbone (Fig. 2B). We found a significant spatial correlation between the nodal CT maturation extent and the mean of its directly connected neighbors (Fig. 2C, adjusted *r* = 0.74, *P* = 5.56×10^-176^). Next, we tested this spatial correlation against two baseline null models. The first model evaluated whether the observed correlation was determined by the wiring topology rather than the basic spatial embedding of the WM network (*44*). Specifically, we generated 1000 surrogate networks by randomly rewiring edges while preserving the nodal degree and approximate edge length distribution of the empirical WM network (“rewired”). The second model evaluated whether the observed correlation was induced by the regional correspondence rather than the spatial autocorrelation of CT maturation (*38, 39*). Specifically, we generated 1000 surrogate maps by rotating region-level cortical *t*-values (“spin test”). After recalculating the correlation coefficient, we found that the observed correlation was significantly higher than the correlations in both null models, and these results were highly consistent for all three nodal resolutions (all *p_rewired_* < 0.001 and all *p_spin_* < 0.001, Fig. 2D). Interestingly, when estimating the spatial constraints at the system level, we found that direct WM connections within the heteromodal area, especially within and between FP and DM networks, showed strong constraints on the maturation of CT (Fig. 2E).

Considering that spatially adjacent nodes may intrinsically exhibit similar cortical development trends, we further performed another two confounding analyses to demonstrate that the observed correlation is not determined by the spatial proximity effect. In the first analysis, we excluded all spatially adjoining neighbors and recalculated the mean CT maturation extent of the remaining structurally connected neighbors for each brain region (“excluded”). In the second analysis, we regressed out the effect of nodal mean Euclidean distance to its connected neighbors from the mean CT maturation extent (“regressed”). After re-estimating the empirical correlation coefficient (1000-node: adjusted *r_excluded_* = 0.60 and adjusted *r_regressed_* = 0.74), we repeated two null model tests and found highly consistent results at all three nodal resolutions (all *p_rewired_* < 0.001, *p_spin_* < 0.001, Fig. S3).

To further validate whether the constraints of direct WM connections on regional CT maturation exist throughout 6 to 14 years old, we also treated age as a continuous variable instead of dividing into two groups. We employed a semiparametric generalized additive model (GAM) (*45*) with sex included as covariates and participant as random effects (Supplementary Text, Section 1.4) to fit the maturation curves of nodal CT with age. Significant age effects of nodal CT were consistently found in the dorsolateral prefrontal regions, lateral temporal, and lateral parietal regions (Fig. 3A, *P* < 0.05, Bonferroni corrected) with most brain nodes showing significant cortical thinning during this period. Two representative fitting curves of nodal CT maturation in the prefrontal cortex and inferior parietal cortex were shown in Fig. 3B. Next, we obtained the maturation rates of nodal CT at each age point by calculating the first derivative of the age smooth function. The brain maps of CT maturation rates at three representative ages were shown in Fig. 3C. Last, we calculated the across-node correlation of the rate of CT maturation between a node and its directly connected neighbor nodes and test this correlation against two baseline null models at each integer age point. We found a significant spatial correlation between the nodal CT maturation rate and the mean of its directly connected neighbors across different age points (*r*: ranged from 0.62 to 0.71, all *p_spin_* < 0.001, all *p_rewired_* < 0.001, Fig. 3D).

**Figure 3.**
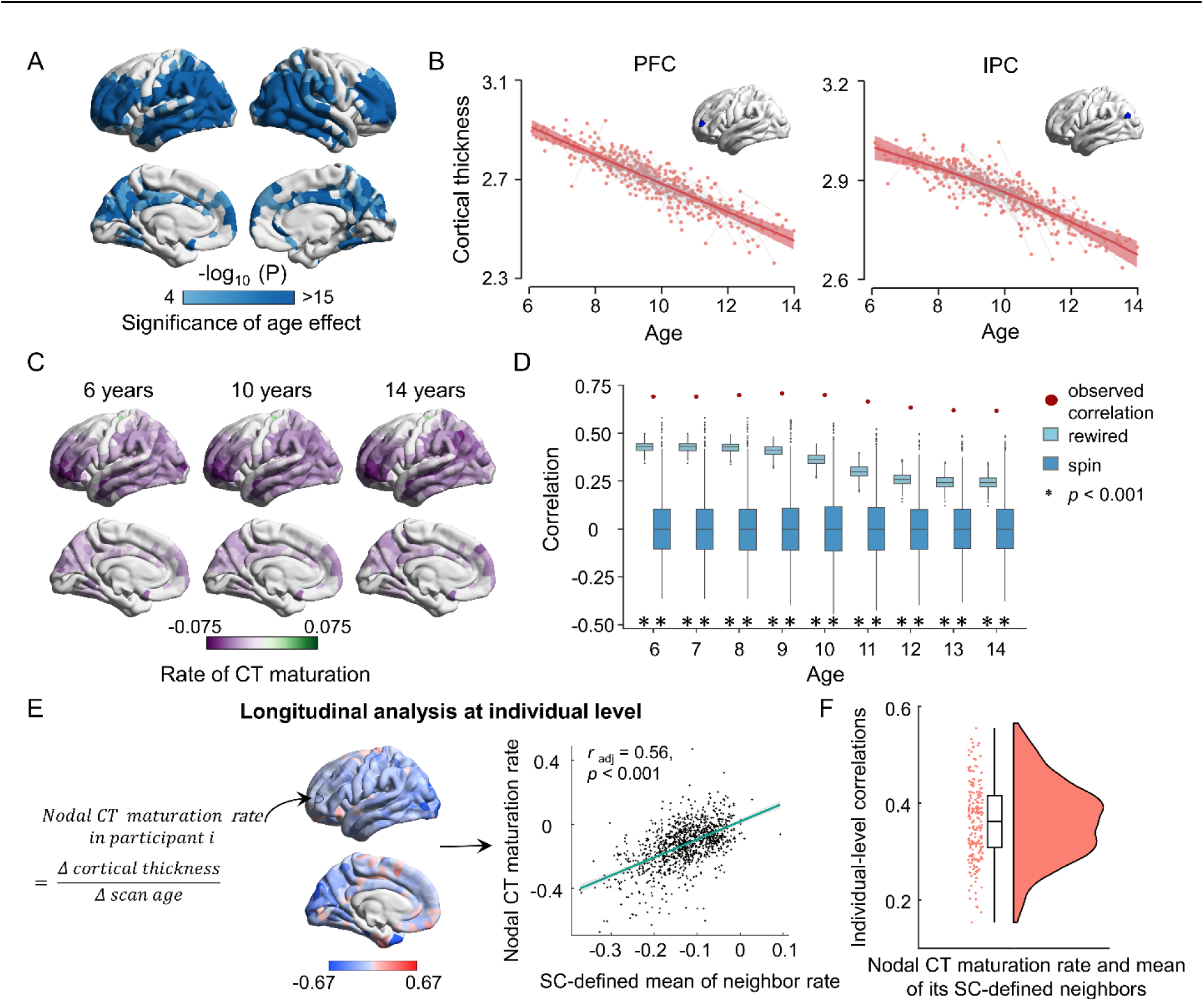
WM network-based CT maturation while either considering age as a continuous variable or performing individual-level analysis. (**A**) Brain map for age effect on the CT maturation from childhood to adolescence using the GAM. The significance levels (-log_10_ P) of the age effect in GAMs are encoded by light to dark blue. The map was corrected using a Bonferroni correction method for multiple comparisons (*p_bonf_* < 0.05). (**B**) Two representative fitting curves of CT maturation in the prefrontal cortex (PFC) and inferior parietal cortex (IPC). (**C**) The brain maps of CT maturation rates (first derivative of the age smooth function) were shown at representative ages of 6, 10, and 14 years. (**D**) Significant correlations were observed between the nodal CT maturation rate and mean of its directly connected neighbors at each integer age point. The observed correlations (red dots) were compared to the correlations obtained from 1000 rewired tests (light blue boxes) and 1000 spin tests (deep blue boxes). Asterisks denote statistical significance (*p* < 0.001). (**E**) Longitudinal WM network-based CT maturation analysis at individual level. Nodal CT maturation rate (mm/year) for each individual was defined as the CT difference between two scans divided by the gap of scan age (ΔCT / Δscan age). Negative values indicate cortical thinning while positive values indicate cortical thickening with development. The brain map of nodal CT maturation rates and the spatial correlation between nodal CT maturation rate and the mean of its SC-defined neighbors for a representative participant was given. (**F**) The distribution of these spatial correlations across individuals. These correlations are significant for almost all participants (*p_spin_* < 0.05 in all longitudinal samples and *p_rewired_* < 0.05 in 98.6% (204/207) of longitudinal samples).

To further assess whether the WM network-constrained CT maturation exists at the individual level, we also utilized all individual longitudinal scans independently (105 participants underwent two scans and 51 participants underwent three scans). We first estimated the brain map of nodal CT maturation rates for each individual (Fig. 3E and Supplementary Text, Section 1.4) by calculating the nodal CT difference between two brain scans divided by the gap of scan ages (ΔCT / Δscan age, Fig. 3E left panels). Next, we reconstrued the individual WM networks and repeated the correlation analysis between nodal CT maturation rates and the mean of its directly WM-connected neighbors within each individual (Fig. 3E right panels). We found that this correlation is significant for almost all individuals (*r*: ranged from 0.15 to 0.56, *p_spin_* < 0.05 in all longitudinal samples and *p_rewired_* < 0.05 in 98.6% (204/207) of longitudinal samples, Fig. 3F).

Collectively, these results provided strong evidence from a network level that the spatial pattern of nodal CT maturation is structurally constrained by the underlying WM network topology.

### The Diffusion Model of the WM Connectome Predicts the Spatial Maturation of CT

To further understand the mechanisms of how the maturation process of cortical morphology is constrained by the WM connectome, we proposed a graph-based diffusion model to simulate the network-level axonal interactions during cortical development. The nodal diffusion processes through multiscale WM edge paths are used to predict the maturation of cortical CT (Supplementary Text, Section 2.4). Specifically, we first calculated the diffusive probabilities of a given node to other nodes during a random walk modeling with *nth* moving steps (for a toy, see Fig. 4A) to represent the nodal diffusion profile at *nth* neighboring scales (*n* = 1, 2, 3, …*N*; the maximum neighboring scales *N* was set as the network diameter, which is the max shortest path length). Increasing moving steps present expansion scales of the probed neighborhood, which indicates local to distributed preferences of information exchange during the diffusion process. The diffusion profiles of all brain nodes form a diffusive probability matrix that represents the distribution of information propagation throughout the whole network. To further characterize the spatial layout of each diffusive matrix, we classified all cortical nodes into seven brain communities (*37*) and calculated the average diffusive probabilities within the same system and between different systems separately across brain nodes. Of note, we observed that the diffusion probabilities within the same cortical system were greater than 0.5 at the 1st scale and then decreasing with the expanding of neighboring scales (Fig. 4B). This indicates that a lower scale is mainly involved in more community segregation during propagation. Then, we trained a support vector regression (SVR) model with nodal diffusive profiles at each neighboring scale separately as input features to predict the CT maturation extent in a 10-fold cross-validation strategy (*46*). To evaluate the significance of prediction accuracy, we compared the empirical accuracy with two null model tests, including a spin test and a rewiring test. We found that the diffusive profiles of a given node could significantly predict its CT maturation extent at multiple neighboring scales (*r_1-9 scale_*: ranged from 0.65 to 0.75, all *p_spin_* <= 0.001, all *p_rewired_* < 0.001, Fig. 4C and Table S1). The prediction accuracies are higher at lower neighboring scales. Additionally, features with high contribution to these predictions are mainly involved in the diffusion of frontal and parietal nodes (Fig. S4). These results are highly consistent across all three nodal resolutions (Fig. S5 and Tables S2-S3). Overall, our analysis of computational models indicates that the diffusive characteristics of WM connectome at the local to distance scales largely determine the spatial maturation map of CT from childhood to adolescence, with a relatively high effect among nodes within the same cortical system.

**Figure 4.**
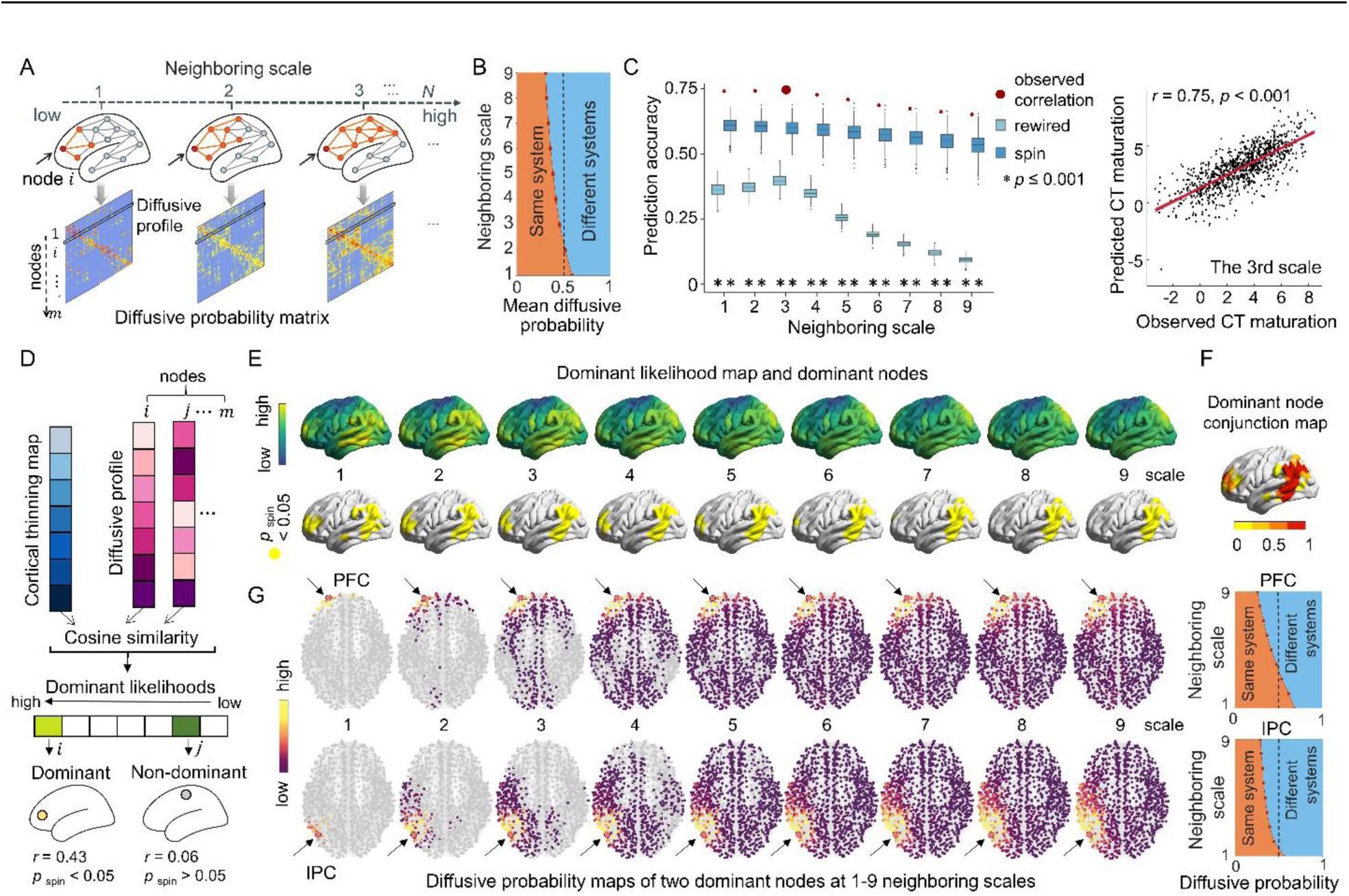
Network-based diffusion model. (**A**) Schematic diagram of nodal diffusion processes through multiscale WM edge paths. The orange color represents the edges and nodes at the *nth* neighboring scale of a given node *i* (red). For the first neighboring scale, the diffusion probabilities of node *i* to its neighbors during one random walk step are 1⁄*d*_*j*_, where *d*_*j*_ is the node degree of *i*. As the moving steps increase, the scale of the probed neighbor nodes also expands and the diffusion probabilities to each neighbor are recalculated iteratively. The diffusive profiles of all nodes form the diffusive probability matrix at each neighboring scale. (**B**) The curve of the average diffusive probability of whole-brain nodes within the same cortical system and between systems. It illustrates that the diffusion probability within the same cortical system was greater than 0.5 at the first scale and then decreases with the expansion of neighboring scales. (**C**) Significant correlations between the predicted CT maturation and the observed CT maturation. The observed correlations (red dots) were compared to the correlations obtained from 1000 rewired tests (light blue boxes) and 1000 spin tests (deep blue boxes). Asterisks denote statistical significance (*p* <= 0.001). The box plot shows the result at 1000-node resolution. See Fig. S5 for results at other resolutions. The scatter plot on the right depicts the correlation between actual and predicted CT maturation at the 3rd neighboring scale, which exhibited the highest prediction accuracy, as an example. (**D**) Schematic of dominant brain region identification. Dominators are regions whose diffusion profiles show significant cosine similarity with the CT maturation map. Spin tests (1000 times) were used to evaluate the statistical significance. (**E**) Regional distributions of dominant likelihood (cosine similarity) between nodal diffusion profiles and CT maturation map at 1-9 neighboring scales (top panels) and the spatial distributions of dominant regions (*p_spin_* < 0.05, bottom panels). Fig. S6A-B and Fig.S7 depict the results in other view directions and in other parcellation resolutions. (**F**) The conjunction map of dominant nodes across all nine neighboring scales shows the probability of each node being identified as a dominant node across scales. (**G**) The diffusive probability distribution of two representative dominant nodes separately in the prefrontal cortex (PFC, top panels) and inferior parietal cortex (IPC, bottom panels) at each 1-9 neighboring scale. A node with a brighter color represents a greater diffusive probability between that node and the dominant node. The right panels show the diffusive probability of both dominant nodes within the same cortical system and between systems. As the neighborhood scale expands, the diffusion of these two nodes spreads from local communities to nearby and distributed communities across whole brain. These diffusion processes were mainly involved in nodes within system at low neighboring scales while in nodes between systems at high scales.

We next try to measure the dominant likelihood map for the spatial constraint between nodal CT maturation and WM connections and screen out brain nodes that lead the whole brain constraint (Supplementary Text, Section 2.5). For each node, we calculated the cosine similarity between its diffusive profiles at the *nth* (*n* = 1, 2, 3, …, *N*) scale and the CT maturation map (Fig. 4D). High similarity of a node indicates that its neighboring diffusion preference largely resembled its neighboring distribution of CT maturation. We observed that the dominant likelihood maps are highly similar across all nine neighboring scales (Fig. 4E, top panel and Fig. S6A) with high values in the bilateral prefrontal, parietal, and temporal regions. These regions were further identified as dominant nodes by higher similarity than expected by chance (*p_spin_* < 0.05) (Fig. 4E, bottom panel, Fig. S6B). The conjunction map of dominant nodes across all neighboring scales is shown in Fig. 4F, where the robust dominant nodes are mainly located in the bilateral prefrontal cortex and inferior parietal cortex. This indicates the leading roles of these regions in shaping the spatial maturation of whole brain CT. Similar results were found in other parcellation resolutions (Fig. S7). To further verify these dominant nodes, we also used a different identification approach (*47*), which defines dominators as brain regions showing high maturation extents in both themselves and their directly connected neighbors (Fig. S8A). Using this approach, we ranked nodes based on their CT maturation extents and their neighbors’ mean CT maturation extents separately in ascending order and then calculated the mean rank of each node across both lists. Regions with higher mean ranks (*p_spin_* < 0.05) were identified as the dominant nodes. We found that this dominant likelihood map was significantly correlated with our network-based diffusion analysis (Spearman’s *r* = 0.81, *p_spin_* < 0.001, Fig. S8B-C), with highly consistent dominant regions (Fig. S8D).

To further exemplify the diffusion processes of the dominant nodes at each neighboring scale, we illustrated the diffusive profiles of the two most robust dominant nodes in the prefrontal region (Fig. 4G, top panels) and inferior parietal region (Fig. 4G, bottom panels), respectively. As the neighborhood scale expands, the diffusion of prefrontal dominators mainly spreads to neighbors within FP and DM systems, while the diffusion of parietal dominators mainly spreads to neighbors within DM, DA, and SM systems (Fig.S6C). These diffusion processes were mainly involved in nodes within the same system at low neighboring scales and in nodes between systems at high neighboring scales.

### Regional Heterogeneous Constraints between CT Maturation and the Connectome Are Associated with Gene Expression Profiles

Next, we sought to explore the genetic associations of the nodal constraints between the spatial maturation of the CT and WM connectomes during development. We adopted the BrainSpan dataset (*48*) (Supplementary Text, Section 3.1), which contains gene expression samples of brain tissues from 8 post-conception weeks to 40 years, to evaluate the regional genetic relevance. We selected four gene sets according to Kang and colleagues (*29*), which cover typical maturation procedures involved in both CT and WM, including axon development, myelination, dendrite development, and synapse development. We hypothesized that there should be differentiated transcriptomic characteristics between the identified dominant and non-dominant brain nodes. To this end, we first divided the cortical tissue samples into two categories according to whether they were dominant nodes in the conjunction map. Then, we calculated the first principal component score of each gene set’s transcription level and estimated the category differences. The statistical significance was calculated by comparing the empirical difference against null differences generated by randomly resampling the same number of genes 1000 times from the remaining genes. We found divergent transcriptomic trajectories between dominant and non-dominant regions in all four maturation processes from childhood to adolescence (Fig. 5A), and the transcription level in dominant regions was significantly higher than that in nondominant regions for dendrite (*P =* 0.014) and synapse development (*P =* 0.002) but significantly lower for axon development (*P <* 0.001) and myelination (*P <* 0.001) (Fig. 5B). This result indicates that gene expression could support the microstructural differences in neurodevelopment between dominant and non-dominant regions, resulting in a non-uniform degree of constraints between CT maturation and WM pathways.

**Figure 5.**
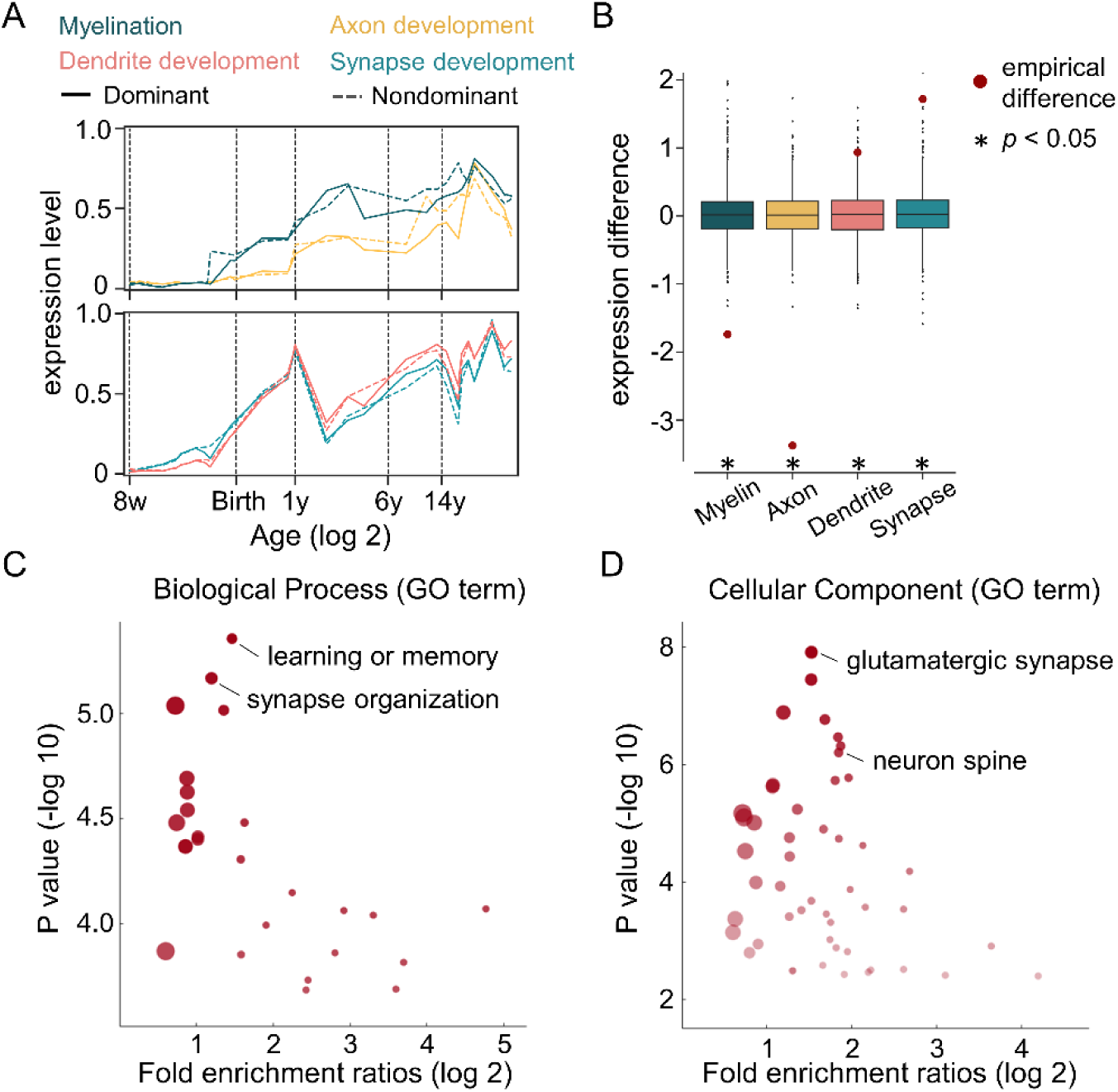
Association between regional heterogeneous constraints and gene expression profiles. (**A**) Transcriptomic trajectories between dominant regions (solid line) and non-dominant regions (dashed line) in four maturation processes. Here, we calculated the first principal component score of each gene set’s transcription level. (**B**) Transcriptomic differences between dominant and non-dominant regions from childhood to adolescence. For each maturation process, the statistical significance was calculated by comparing the empirical difference (red dots) against null differences generated by randomly resampling the same number of genes 1000 times from the remaining genes. Asterisks denote statistical significance (*p* < 0.05). (**C**) Volcano plot depicts Gene Ontology (GO) results for Biological Processes and Cellular Components (**D**). The dots represent the GO terms corrected for multiple comparisons (FDR-corrected, *P* < 0.05). The size of the dot indicates the number of genes belonging to the corresponding GO term, and the transparency of the dot represents the significance of the corresponding GO term.

Considering that the BrainSpan dataset only contains 11 sampling neocortex areas, we also validated the regional gene expression relevance by using Allen Human Brain Atlas datasets (*49*) Supplementary Text, Section 3.2). After preprocessing with the abagen toolbox (*50, 51*), a matrix of gene expression profiles was generated (111 left brain regions × 8631 gene expression levels). Then, we identified the association between the dominant likelihood map and each gene expression map using Pearson’s correlation and spin tests (1000 times). A total of 457 genes showed a positive correlation, and 619 genes showed a negative correlation (*p_spin_ <* 0.05, FDR corrected, Table S6). Next, we performed Gene Ontology enrichment analysis (Supplementary Text, Section 3.3) on these two gene sets using the ToppGene Suite (*52*). We found a significantly correlated gene list with positive correlations mainly enriched in learning or memory and synapse organization (biological process) as well as glutamatergic synapse, neuron spine, and somatodendritic compartment (cellular component) (all *P <* 0.05, FDR corrected, Fig. 5C, Fig. 5D) and negative correlations enriched in the generation of precursor metabolites and energy process (biological process) and myelin sheath components (cellular component) (all *P <* 0.05, FDR corrected, Fig. S9). The detailed enrichment analysis results are shown in Tables S7-S8.

### Sensitivity and Replication Analyses

Considering that the diffusion weighting scheme of Discovery dataset only includes b-values of 1000, which could affect the effectiveness of the WM tractography (*53*), we used another independent diffusion imaging dataset with multi-shell diffusion gradients that contain high b-values shells from HCP-D (*32*) to reconstruct the individual WM network and regenerate the group backbone. Using this new backbone, we found highly consistent results with our main findings. Specifically, nodal CT maturation extents are significantly correlated with their directly connected neighbors (adjusted *r* = 0.76, *p_spin_* < 0.001, *p_rewired_* < 0.001, Fig. S10A). Using the network-based diffusion model, the spatial maturation of CT was also predicted by the diffusion properties of the WM network (*r_1-8 scale_*: ranged from 0.69 to 0.78, all *p_spin_* < 0.001, *p_rewired_* < 0.001, Fig. S10B and Table S4).

To evaluate the reproducibility of our findings, we further replicated all main analyses using the multi-site Replication dataset from HCP-D. The site item was set as a random effect in both linear mixed models and GAM models to control for site effects. The results are highly consistent with those obtained using the Discovery dataset: (i) several heteromodal areas including dorsolateral prefrontal regions and lateral parietal regions exhibited the most pronounced cortical thinning (Fig. S11A); (ii) CT maturation extent (*t*-value) of a node shows a positive correlation with the mean maturation extent of its directly connected neighbors (adjusted *r* = 0.62, adjusted *r_excluded_* = 0.46, and adjusted *r_regressed_* = 0.62), and the empirical correlation exceeded the values in null models (all *p_spin_* < 0.001 and all *p_rewired_* < 0.001, Fig. S11B-D); (iii) nodal CT maturation rate is significantly correlated with the mean of its directly connected neighbors at each age point (*r*: ranged from 0.41 to 0.66, all *p_spin_* < 0.001 and all *p_rewired_* < 0.001, Fig. S11E); (iv) the diffusion profiles of the WM network at multiple neighboring scales also predicted the spatial maturation of CT (*r_1-4 scale_*: ranged from 0.60 to 0.66, all *p_spin_* < 0.05, *p_rewired_* < 0.001, Fig. S11F and Table S5); and (v) dominant nodes mainly reside in the lateral parietal regions (Fig. S11G). Taken together, these findings provide replicable evidence that the WM network constrains the spatial maturation of CT from childhood to adolescence.

## Discussion

The present study shows for the first time the constraints of WM network architecture on the coordinated maturation of regional CT from childhood to adolescence and their associations with gene expression signatures. Specifically, we proposed a network-based diffusion model to predict regional cortical maturation from WM connectome architecture. These constraints are regionally heterogeneous and regulated by the gene expression of microstructural developmental processes. These results are largely consistent across three cortical parcellations and are highly reproducible across two independent datasets. Taken together, these findings provide insights into the understanding of network-level mechanisms that support the maturational process of cortical morphology.

Numerous previous studies have documented that the human brain undergoes remarkable refinements during childhood and adolescence, such as cortical thinning, area expansion, and WM myelination (*2, 4, 7, 54, 55*). These multifaced gray matter and WM changes have been proven to be intrinsically linked with each other at the regional level. For example, in early childhood, the spatial pattern of cortical surface area expansion during development is highly similar to the myelination of underlying cortico-cortical tracts (*56*). In children and adolescents, Jeon et al. (*19*) reported a significant correlation between the rate of CT decrease and the rate of FA increase in WM tracts at local gyri of the frontal lobe. Ball et al. (*57*) observed a shared developmental process in CT and structural connectivity during childhood and adolescence. In addition, prior studies show that homologous cortical regions tightly connected by rich WM tracts show high CT maturation couplings (*21, 23*). At the microscale level, cortical morphology changes during maturation are thought to have various biological origins, including synaptic pruning, increased axon diameter, and myelination (*54, 58*). Seeking a unified original model for the whole-brain cortical changes is difficult since even within the ventral temporal cortex, thinning of different brain regions seems to be due to distinguished factors (*58*). Here, we address this issue with a new perspective on brain network modeling. We showed that the morphology maturation of cortical nodes is well represented by that of their WM-connected neighbors during the transition from childhood to adolescence even after excluding the spatial proximity effect (Fig. 2B-D, Fig. S3). It was highly reproducible at different age points, within individuals, and in an independent dataset (Fig. 3D and Fig. 3F, Fig. S11). Such a network-level association is an important extension of previous developmental theories that support cortical thinning across brain development.

The WM network-based cortical maturation could be explained by several factors as follows. First, animal studies revealed that cortical regions that are structurally connected by axon projections are more likely sharing similar cytoarchitectures, such as neuronal density and laminar differentiation (*59, 60*). Moreover, higher cytoarchitectural similarity among regions tend to higher cortical coordinated maturations (*61, 62*) among neighboring nodes in the brain WM network. Second, a recent study using 19 different neurotransmitter receptors/transporters, such as dopamine and glutamate, found that structurally connected cortical regions usually show greater neurotransmitter receptor similarity (*63*). Therefore, these regions may be more inclined to be coregulated by similar physiological processes during development (*64, 65*). Third, direct WM connections facilitate ongoing interregional communication, enabling these regions to exhibit strong spontaneous neuronal activity couplings (*66*), which indicates the natural preference for the regional coordination of functional development. This also coincides with Hebbian learning rule, where neurons that fire synchronously tend to form or consolidate connections between them (*67, 68*). Additionally, WM network-based constraints on cortical morphology exist extensively in adult brains. For instance, Gong et al. suggested that approximately 40% of edges in the adult CT covariance network show matched WM connections (*69*). In degenerative brain diseases, including schizophrenia, dementia, and Parkinson’s disease, studies also found that the disease-related cortical deformation pattern across brain regions is conditioned by the WM network (*47, 70, 71*).

Notably, in this study, we proposed a network-based diffusive model for the constraint of WM on CT maturation. We highlight that nodal diffusion profiles of the WM connectome could accurately predict the maturation pattern of regional CT (Fig. 4C). From a physical transport perspective, the axonal microenvironment can be regarded as a porous medium that makes diffusion processes within brain tissues extremely critical for delivering oxygen and glucose during neuron metabolism (*72*). Meanwhile, diffusion of chemical neurotransmitters at synaptic clefts along axons is essential for forming postsynaptic responses during intercellular communications (*73*). At the macroscopic scale, network-based models have been proposed to simulate the consequences of interregional diffusive spread in latent topological space throughout the brain connectome. In neurodegenerative diseases (e.g., Alzheimer’s disease), these models showed excellent prediction abilities for the spatial atrophy pattern of the cortex by capturing disrupted transport of trophic factors or accumulated spread of toxic misfolded proteins (*74, 75*). Based on brain images of nine very prematurely born infants, Friedrichs-Maeder et al. employed a diffusion model to explore the relationship between WM connectivities and cerebral MR measurements such as T1 relaxation time (*76*). They reported that early maturation in the primary sensory cortex serves as a source to gradually propagate into the higher-order cortex. In our study, considering the intricate biological relevance between brain WM and cortical morphology (*54, 58*), we used a simple random walk model to depict the complex network diffusive processes of brain nodes. This model can concisely present the local to distributed supports of the structural connectome on cortical maturation from childhood to adolescence. These nodal diffusive features are effectively integrated by a multivariable machine learning model to represent nodal cortical maturation. Of note, this model first showed the significance of indirectly WM-connected neighbors for constraining nodal morphology maturation, which strongly emphasizes the necessity of employing a network-level model to capture this relationship. The contribution from indirect neighboring scales is reasonable because cortical communications between brain regions inherently contain high-order components to support information exchanges between topologically distant nodes (*77, 78*). Meanwhile, these indirect WM neighbors are shown majorly located within nearby cortical communities (Fig.4B-C) that share common maturation processes to support morphological integration during cortical development (*79, 80*).

Our results also showed that the constraints of the WM network on CT maturation are spatially heterogeneous (Fig. 4E). Regionally, dominant nodes in the heteromodal area, especially within and between FP and DM networks, show the strongest spatial constraints. Previous neuroimaging studies have revealed that FP and DM networks display dramatic cortical thinning from childhood to adolescence (*1, 8*). During the same period, brain WM fractional anisotropy and functional connectivity also show prominently increased tendencies within these networks (*81, 82*). Our results imply that the strong WM constraints on the cortical maturation of the heteromodal area may determine the major pattern of whole-brain cortical thinning. Compatible with our findings of connectome-morphology constraints, structure‒function association studies also show age-related increases in heteromodal area during youth, which are associated with individual executive performance (*83*). These multifaced heteromodal refinements could support the rapid enhancement of high-order cognitive and social capabilities such as working memory and reasoning (*82, 84*).

By employing transcriptome imaging analyses in a developmental gene expression dataset, we found that those dominant nodes in the heteromodal area show different transcriptional patterns compared with non-dominant brain nodes. Specifically, dominant regions exhibit higher gene expression levels involved in the maturation of gray matter morphology, including synaptic and dendritic development, and lower expression levels associated with WM maturation, such as axon and myelin development. This coincides with the findings from histological samples and MR images studies, demonstrating that heteromodal regions have higher synaptic density and lighter myelination than other regions in childhood and adolescence. This brings prolonged maturation of high-order cortex during adolescence to support the optimization and consolidation of synaptic and axonal connectivity compatible with cognitive growth and the environment (*1, 26, 28, 55*). Likewise, we conducted GO enrichment analysis with the AHBA dataset, which is the most complete gene expression dataset available on the human brain to date, and found that the nonuniform degree of constraints is mainly related to the biological processes and cellular components involved in learning or memory, synapse organization, glutamatergic synapse, and neuron spine. These gene-related processes are involved in the spatial thinning of CT during childhood and adolescence (*26, 27*). As the most abundant synapse type in the neocortex, glutamatergic synapses are primarily responsible for the transport of excitatory transmitters, which are crucial for regulating the transmission and processing of information among brain regions (*73*). Meanwhile, neuron spines on dendrites serve to receive various kinds of excitatory inputs from axons and are considered crucial for brain circuit wiring distribution and circuit plasticity (*85, 86*). Disruptions of these synapses structures are important substrates of pathogenesis in multiple neurodevelopment diseases, especially those with deficits in information processing, such as autism (*85, 87*). In summary, our findings provide evidence that genetic factors associated with microstructure development contribute to these connectome-based constraints on cortical maturation.

Several issues need further consideration. First, although dMRI-based deterministic tractography used here is currently a common approach for reconstructing WM tracts in vivo, it is still inherently limited, especially for depicting cross-fibers (*88, 89*). Although probabilistic tractography approaches exhibit high sensitivity for this issue (*90*), they result in excessive false positive connections (*89*). Second, diffusive processes during axonal transport are proven directional (*13*). However, *in vivo* inference for the direction of WM fibers is still extremely difficult with tractography-based methods. Future investigations combining diffusion models with animal connectome by molecular tracers would reveal a directed network constraint mechanism. Third, dMRI is an indirect way to gain sensitivity to WM microstructure. There still exists many limitations in characterizing intra-axonal properties, particularly at lower diffusion weightings. Thus, in the present study we employed the binary networks to capture the backbone of the WM connectome. In the future, advanced imaging techniques such as quantitative MRI could be utilized to better capture the microstructural properties of brain tissues and further understand the relationship between WM network development and cortical morphological maturation. Fourth, the developmental gene data from BrainSpan only contain 11 areas of the neocortex (*48*), which can only provide a rough exploration of the differences in gene expression between dominant and non-dominant nodes. We further validated this result using the AHBA datasets (*49*), but it was sampled from only six postmortem adult brains. Future studies with gene expression data of widespread cortical regions from a large sample of children and adolescents would be important for connectome-transcriptome association analysis. Finally, we showed the constraints of the WM network on cortical morphology maturation in typical development. Previous studies have documented both abnormal cortical maturation and WM connectome structure in neurodevelopmental disorders such as autism (*91*) and attention-deficit/hyperactivity disorder (*92*). In the future, it would be desirable to examine how the WM connectome shapes cortical morphology in these atypical populations.

## Materials and Methods

### Participants and Data Acquisition

We performed analyses in two independent datasets. After quality control, the Discovery Dataset included a longitudinal cohort of 314 participants (aged 6-14 years) with 299 scans in the child group (mean (± SD) age 8.68 ± 0.94 y, 157 females) and 222 scans in the adolescent group (mean age 11.19 ± 0.94 y, 108 females) from the Beijing Cohort in Children Brain Development (CBD) project (*93*). Among all participants, 158 underwent a single scan, 105 underwent two scans with a mean time interval of 1.16 years, and 51 underwent three scans with a mean time interval of 0.99 years. Structural and high angular resolution diffusion imaging (HARDI) diffusion MR brain images for each subject were scanned at Peking University using a 3T Siemens Prisma scanner. Informed written consent was obtained from all participants and at least one parent/guardian, consistent with the guidelines of the Ethics Committee of Beijing Normal University. The Replication Dataset included a cross-sectional cohort of 301 participants (aged 5-14 years) with 98 scans in the child group (mean age 8.72 ± 0.99 y, 32 females) and 203 scans in the adolescent group (mean age 12.17 ± 1.28 y, 86 females) selected from the Lifespan Human Connectome Project in Development (HCP-D) (*32*). Participants were recruited across four imaging sites, and details on imaging protocols can be found in (*94*).

### MRI Data Preprocessing

For the Discovery Dataset, cortical reconstruction was performed using FreeSurfer v6.0 image analysis suite (https://surfer.nmr.mgh.harvard.edu/). This processing includes intensity normalization, nonbrain tissue removal, tissue segmentation, automated cortical reconstruction, and surface parcellation (*34, 95–98*). To reconstruct the individual cortical surface, all images were first processed cross-sectionally and then processed through the longitudinal stream (*99, 100*) in FreeSurfer to obtain more sensitive and reliable measurements of cortical morphology. Next, we constructed a custom registration template by averaging all available subjects’ cortical surfaces. The atlas in the standard *fsaverage* space was registered to the new custom template and then registered to each subject’s surface space to be used to obtain regional CT measurements. All images were visually inspected and manually edited and corrected where needed to ensure the correctness of gray matter and WM boundaries and improve the quality of the output. Diffusion data were first denoised, and Gibbs ringing artifacts (*101*) were removed using MRtrix 3.0 (*102*). Next, we corrected eddy current-induced distortions, head movements, and signal dropout using the FSL eddy tool (*103–105*). Then, to make the susceptibility-induced EPI distortion correction, we fed the eddy-corrected DW images and corresponding fieldmap images into the FUGUE tool (https://fsl.fmrib.ox.ac.uk/fsl/fslwiki/FUGUE/Guide#Making_Fieldmap_Images_for_FEAT) to remove EPI susceptibility artifacts. Finally, B1 field inhomogeneity was corrected for the dMRI images with the N4 algorithm available in ANTS (*106*).

For the Replication Dataset, the T1w data went through the HCP preprocessing pipeline (*107*). We obtained the individual CT in a common 32k_fs_LR space from the publicly available dataset. Diffusion data were first denoised, and Gibbs ringing artifacts were removed. Then, we used topup/eddy (*103–105, 108*) to correct the EPI distortions, eddy currents, subject movement distortions, and signal dropout. Finally, B1 field inhomogeneity was corrected for the dMRI images.

### Estimation of Regional CT and WM Networks

Each participant’s cortex was parcellated into 1000 regional nodes with approximately equally sized based on the modified Desikan-Kiliany atlas (*34, 35*) and verified at 219-node and 448-node parcellations. The CT of each brain node was estimated by using FreeSurfer v6.0 software (https://surfer.nmr.mgh.harvard.edu/). Then, we reconstructed the whole brain anatomical streamlines using native diffusion MR images for each individual by employing generalized q-sampling imaging (GQI)-based deterministic streamline tractography (*42, 109*) with gray-white boundary as seed voxels. Two cortical regions were considered structurally connected if there exists at least one streamline with two end points located separately in them (*110, 111*). After obtaining the individual WM network, we further implemented a consensus approach to generate the binary group-level WM connectome (*43*).

### Analysis of CT Maturation from Childhood to Adolescence

We explored the CT maturation by the following analysis. (i) To estimate the maturation of CT from childhood to adolescence, we applied a mixed linear analysis with the sex term included as the covariate and the group term as the main effect for each brain node. The model was defined as follows:

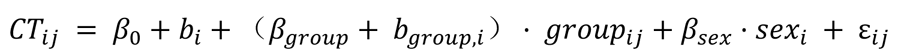

Specifically, *CT*_*ij*_ is the CT of participant *i jth* scan, *β*_*group*_ represents the fixed group effect of participant *i*, *b*_*group*_ is the random effect, and ε_*ij*_ is the residual. The *T* statistics from the group term were used to represent the maturation extent of brain nodes. Greater positive *t*-values indicated more significant cortical thinning. (ii) To further validate whether the constraints of the WM network architecture on CT maturation exist throughout 6 to 14 years old, we considered age as a continuous variable using the semiparametric generalized additive models (GAM) (*45*) to flexibly investigate linear and non-linear relationships between CT and age. For each cortical node, the model was defined as follows:

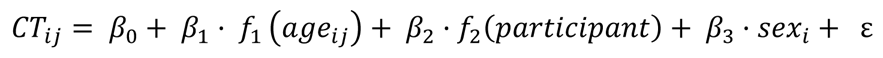

where CT of participant *i* at the *jth* scan as the dependent variable, age as a smooth term, sex as linear covariates and participant as random effects. Thin plate regression splines were used for the smoothing basis and the residual estimates of maximum likelihood (REML) method was used to estimate the smoothing parameter. Next, we calculated the first derivative of the age smooth function (Δ cortical thickness / Δ scan age) to characterize the CT maturation rate. (iii) To assess whether the WM network-constrained CT maturation also exists at the individual level, we estimated the individual-level CT maturation rates using the longitudinal MRI scan data from each participant in the Discovery Dataset. The participant who underwent three repeated scans was split into two continuous pair-scan combinations. Therefore, a total of 207 longitudinal samples were included in this analysis. For each brain node, the CT maturation rate was defined as follows:

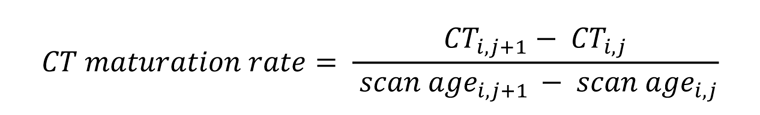

where *CT*_*ij*_ is the CT of participant *i* at the *jth* scan. Negative values indicate cortical thinning while positive values indicate cortical thickening with development.

### Association between CT Maturation and WM Connectome

To test whether the regional maturation of CT was constrained by its direct WM connections, we first assessed the across-node relationship between the CT maturation extent (*t*-value between child and adolescent groups) of a node and its directly connected neighbor nodes by a model as follows:

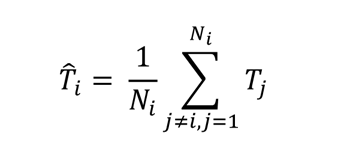

In this model, *T*_*i*_ represents the predicted CT maturation extent of node *i* according to its directly connected neighbors. *T*_*j*_ represents the CT maturation extent (*t* values as mentioned above) of the *jth* neighbor, and *N*_*i*_ is the number of directly connected neighbors of node *i*. Specifically, we used the group-level binary WM network to define the WM-connected neighbors of each cortical node. Then, we calculated the spatial correlation between the empirical CT maturation extent (nodal *t*-value) and the predicted values (^*T*^^_*i*_). The correlation coefficient was used to represent the constraint degree between the WM edges and the nodal maturation pattern of CT. We conducted a similar analytical process to quantify the constraint degree for each year of CT maturation rate obtained through GAMs.

To conduct individual level analysis, we reconstructed the individual WM network for each participant from their first scan in each pair-scan combination. Then, we calculated the spatial correlation between the nodal CT maturation rate and the mean of its directly connected neighbors in individual WM network

### Null Models

We tested the observed spatial correlation against two baseline null models. In the first null model, we used a spatial permutation test (“spin test”) to explore whether the observed correlation is specific to the actual CT maturation pattern rather than the spatial autocorrelation of CT maturation (*38, 39*). Specifically, we first record the spherical coordinates of centroids for each parcel in the Cammoun atlas (*35*). Then, we randomly rotated the parcels while maintaining spatial autocorrelation and reassigned node values to the nearest parcels. This procedure was repeated 1000 times to create surrogate brain maps. The *p*-value was calculated as the fraction of correlations in null models exceeding the observed correlation.

In the second null model, we evaluated whether the observed correlation is determined by the empirical WM network topology rather than the basic spatial embedding of the WM network (such as the distribution of node degree and edge length), we used a rewired null model (“rewired”) (*44*). Specifically, we first divided edges into different bins according to their Euclidean distance. To preserve the degree sequence and approximate edge length distribution of the empirical WM network, edge pairs were randomly swapped within each bin. Finally, 1000 surrogate networks were generated by repeating this procedure. The *p*-value was calculated as the fraction of correlations in null models exceeding the observed correlation.

### Network-based Diffusion Model

We proposed a diffusion model by combining *nth*-order random walk processes with an SVR method to determine whether the diffusion properties of the WM network could predict the maturation pattern of CT. Specifically, for an adjacency matrix *A*, the probability of node *i* transferring to its neighbor *j* during one step is *A*_*ij*_/*d*_*j*_ (modeled by a random walker moving one step along the edges of the WM network), where *d*_*i*_ is the structurally connected neighbor number (node degree) of node *i*. Thus, the transition probabilities of the WM network were represented by the transition matrix P. P was defined as:

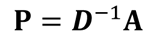

where ***D*** is the node degree diagonal matrix. The initial distribution of random walkers is represented in ***p***_0_, where the diagonal elements are 1 and the other values are equal to 0. Therefore, when these random walkers move *n* steps (*n* = 1, 2, 3, …), their distribution can be described as:

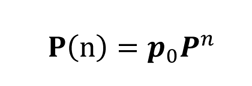

The sum of elements in each row of the distribution matrix is 1, reflecting the diffusion preference of each node with its *nth*-order neighborhoods. Finally, we averaged the outgoing and incoming random walker distribution matrix as a symmetrical diffusion connectivity matrix at each step to represent the bidirectional diffusion properties between any two nodes. Each row of this matrix represents the diffusive profile at the *nth* neighboring scale of each cortical node.

Next, we trained an SVR model with diffusive profiles at all neighboring scales of a brain node as input features to predict its nodal CT maturation extent. This model was trained in a 10-fold cross-validation strategy with a linear kernel. The Pearson correlation coefficient between the empirical and predicted CT maturation extents was calculated as the prediction accuracy. To evaluate the contribution of each scale, we also use the diffusive profiles at each neighboring scale as the input features to further verify the results. Two null models were used to evaluate the significance of prediction accuracy (see Null Models).

### Identifying the Dominant Regions during Development

To further identify the dominant regions, which play more important roles in leading cortical development, we calculated the cosine similarity between the CT maturation map and the nodal diffusion profiles at each random walk step. The statistical significance of the spatial similarity for each brain region was assessed by using spin test (1000 times, see Null Models). Regions with significantly greater spatial similarity (*p _spin_* < 0.05) were identified as the dominant regions during development.

We further replicated our results using the other method introduced by (*47*), which aims to find some brain regions that show high maturation extents in both themselves and their directly connected neighbors. To identify such regions, we ranked the nodes’ CT maturation extents and their neighbors’ mean CT maturation extents in ascending order. For each node, we calculated the mean rank across both lists. Regions with significantly higher ranks (*p _spin_* < 0.05) were identified as the dominant regions.

### Analysis Relationship between Heterogeneous Connectome Constraints on Cortical Maturation and Gene Expression Profiles

We used developmental gene expression data from BrainSpan (*48*) to evaluate whether there are distinctions in the expression levels of genes associated with several neural development events between dominant and non-dominant regions. We divided tissue samples into dominant and non-dominant categories according to their anatomical location (from 11 areas of the neocortex) and arranged them in ascending order based on age to explore the temporal characteristics of gene expression. Next, four typical maturation gene sets (*29*) were selected covering axon development, myelination, dendrite development, and synapse development to evaluate whether there are differences in transcription levels between dominant and non-dominant regions. For each gene set, we performed principal component analysis (PCA) on the gene expression matrix to calculate the first principal component score of each gene set’s transcription level in dominant and non-dominant regions. Then, we calculated the difference between the means of the first principal component scores of the two categories of brain regions. Finally, we randomly sampled an equal number of genes with each gene set from the remaining genes in BrainSpan datasets and recalculated the difference and compared the observed transcription level differences against the null distributions generated by repeating 1000 permutation tests. To further validate the relationship between spatial heterogeneous constraints and cortical gene expression levels at the whole-brain level, we performed a Pearson’s correlation analysis with Allen Human Brain Atlas datasets (*49*) combined with Gene Ontology enrichment analysis.

Detailed information about participants, image acquisition, data preprocessing, and data analyses are further described in Supplementary Materials.

## Acknowledgments

This work was supported by the National Key Research and Development Project (No. 2018YFA0701402), the National Natural Science Foundation of China (Nos. 82021004, 31830034, 31521063, 31221003), Changjiang Scholar Professorship Award (No. T2015027), the Beijing Brain Initiative of Beijing Municipal Science & Technology Commission (No. Z181100001518003) and the China Postdoctoral Science Foundation (202050 and 2022M710433). We thank Dr. Yongbin Wei for the discussion on gene expression data. We thank the National Center for Protein Sciences at Peking University in Beijing, China, for assistance with MRI data acquisition. We also thank the Allen Institute for Brain Science for providing the gene expression data. Research reported in this publication was supported by the National Institute of Mental Health of the National Institutes of Health under Award Number U01MH109589 and by funds provided by the McDonnell Center for Systems Neuroscience at Washington University in St. Louis. The content is solely the responsibility of the authors and does not necessarily represent the official views of the National Institutes of Health.

## Author contributions

X.Y.L., T.D.Z., and Y.H. designed research; W.W.M., Y.P.W., S.P.T., J.H.G., S.Z.Q., S.T., and Q.D. collected the imaging dataset; T.D.Z., L.L.S., X.H.L., T.Y.L., M.R.X., D.N.D., Z.L.Z., Q.L.L, Z.L.X., and Y.H. provided the methodological instruction; X.Y.L. and T.D.Z. performed the data analysis; X.Y.L., T.D.Z., and Y.H. wrote the paper; X.Y.L., L.L.S., T.D.Z., and Y.H. revised the paper.

## Data and materials availability

The Replication Dataset used here is from the Lifespan Human Connectome Project in Development (*32*), which is available for download through https://nda.nih.gov/. The BrainSpan Atlas dataset is available at http://brainspan.org/static/download.html (*48*). The AHBA dataset is available at https://human.brain-map.org/static/download (*49*). The study data and codes are available at https://github.com/Xinyuan-Liang/SC-shapes-the-maturation-of-cortical-morphology.

